# Light effects on circadian and homeostatic regulation: alertness increases independent of time awake

**DOI:** 10.1101/2020.04.17.047423

**Authors:** R. Lok, T. Woelders, M.J. van Koningsveld, K. Oberman, S.G. Fuhler, D.G.M. Beersma, R.A. Hut

## Abstract

Light induced improvements in alertness are more prominent during night-time than during the day, indicating circadian regulation or wake duration related dependence. Relative contributions of both factors can be quantified using a forced desynchrony (FD) designs. Here we investigate alerting effects of light in a novel 4×18 hours FD protocol (5h sleep, 13h wake) under dim (6 mlux) and bright light (1159 mlux) conditions. Hourly saliva samples (melatonin and cortisol assessment) and 2-hourly test-sessions were used to assess effects of bright light on subjective and objective alertness (electroencephalography and performance). Results reveal (1) stable free-running cortisol rhythms with uniform phase progression under both light conditions, indicating that FD designs can be conducted under high intensity lighting, (2) subjective alerting effects of light depend on elapsed time awake, while (3) light consistently improves objective alertness independent of time awake. Three dimensional graphs reflecting light induced alertness improvements depending on wake duration related variation and circadian clock phase suggest that performance is improved during daytime, while subjective alertness remains unchanged. This suggests that light during office hours might be beneficial for performance, even though this may not be perceived as such.

## Introduction

The suprachiasmatic nucleus (SCN) is the pacemaker of the mammalian circadian timing system and regulates daily cycles in activity, hormonal levels and other physiological variables ^1,2^. The primary input of the SCN is light, and this pathway possibly plays an important role in acute alerting effects of light ^3^.

A high level of alertness is defined as a state of high sensitivity to incoming stimuli ^4^. It is known to affect many psychological and physiological functions, such as performance, caloric intake and pain sensitivity ^5–8^. Polychromatic white light has been shown to significantly improve alertness during the evening and night ^3^. However, effects during daytime are less conclusive (reviewed in ^9^). Together, these studies implicate differences in alerting effects of light at different times of day.

Alertness decreases with wakefulness duration, but also fluctuates independent of time since sleep offset, with a cycle of approximately 24 hours. This indicates an influence of the SCN ^10,11^. The circadian drive for alertness increases during the day when homeostatic sleep-pressure levels increase, resulting in relatively stable levels of alertness during the wake period ^12^. Discrepancies in alerting effects of light reported by studies investigating these effects at different times of day suggest that light effects on alertness might be modulated by circadian clock phase and/or by the amount of accumulated sleep pressure.

One approach to distinguish between circadian- and wake duration related variation is a forced desynchrony (FD) paradigm. In a FD paradigm, the wake periods are scheduled with a period that deviates sufficiently from 24 hours such that it falls outside the range of circadian entrainment by light. This allows the internal pacemaker to free run (i.e. following its intrinsic circadian period) throughout scheduled sleep and wakefulness ^13^. As a consequence, sleep and wake intervals are scheduled at different circadian clock times, resulting in multiple combinations of homeostatic sleep drive levels and circadian clock time along the FD protocol. Under certain assumptions, it is possible to mathematically disentangle homeostatic- and circadian clock phase effects on parameters of interest ^12,14^.

FD protocols have only been performed in dim light (<10 lux). Light is known to phase shift the SCN which can interfere with methods of disentangling wake duration related variation from circadian components. This is partially caused by relative coordination of the circadian system with the sleep-wake pattern and synchronous light-dark cycle. However, examination of the parameter space of the human circadian pacemaker indicated the possibility to run a FD design at various light intensities without major effects of relative coordination ^15^. To investigate the contribution of both circadian and wake duration related variation to alerting effects of light, we conducted such an FD experiment in humans under both low and high light intensities.

## Materials and methods

### Mathematical simulations

Kronauer’s model of the human circadian pacemaker ^15^ was used to model clock phase changes during a 72 hour forced desynchrony protocol, consisting of 4 × 18h (5 hours sleep – 13 hours wakefulness) under dim light (DL; 6 mlux) and bright light (BL; 1159 mlux) conditions. Simulations with intrinsic circadian periods (tau) ranging from 24 to 24.6 hours indicated a free running rhythm of the circadian pacemaker with uniform phase progression under both light conditions (Fig S1). Simulations with the two process model of sleep regulation ^16^ allowed estimation of sleep pressure build-up under different sleep-wake durations. Accumulated sleep pressure at the end of wakefulness after 5 h sleep – 13 h wakefulness was similar to a 8h sleep - 16h wake cycle (Fig. S2). According to the model sleep pressure levels did not systematically increase or decrease over the course of the FD protocol (Fig S2).

### Subjects

Participants were eight healthy, non-sleep deprived males between the ages of 20 and 30 years (average ± SEM; 24.0 ± 1.16). All participants provided written informed consent and received financial compensation for participation. The study protocol, screening questionnaires and consent forms were approved by the medical ethics committee of the University Medical Center Groningen (NL54128.042) and were in agreement with the Declaration of Helsinki (2013).

An in-house developed general health questionnaire was used to assess health of the participants. As an indication of sleep timing, chronotype was assessed via the Munich Chronotype Questionnaire (MCTQ ^17^). To determine baseline sleep quality, participants completed the Pittsburgh Sleep Quality Index (PSQI ^18^).

Participants reported no health problems, were intermediate chronotypes (Midpoint of sleep on work-free days, sleep-corrected (MSF_sc_) average ± SEM; 4.88 ± 0.60) and did not report more than mild sleep problems (PSQI<12, average ± SEM; 4.63 ± 1.13). Exclusion criteria were: 1) chronic medical conditions or the need for (sleep)medication use, 2) shift work within three months before participation, 3) having travelled over multiple time-zones within two months before participation, 4) smoking, 5) moderate to high levels of caffeine intake (more than 4 cups per day), 6) excessive use of alcohol (>3 consumptions per day), 7) use of recreational drugs in the last year, 8) a body mass index outside the range of 18 to 27, 9) inability to complete the Ishihara color blindness test ^19^ without errors upon arrival. The estimated average ± SEM of caffeine intake was 0.75 ± 0.35 cups per day for included participants.

### Protocol

Subjects were equipped with wrist worn Actigraphy (CamNtech, United Kingdom) 5 weeks before the start of the in-lab FD experiment to monitor regularity in sleep/wake cycles. On the first day of the lab protocol (Fig 1), participants arrived at the human isolation facility of the University of Groningen 10 hours before habitual sleep onset (HSon; assessed with the MCTQ). Upon arrival, individuals were equipped with EEG electrodes at the left and right frontal, central and occipital locations, two electro-oculogram (EOG) electrodes placed above and below both eyes, and two electrodes above the eyebrow for electromyography. Two reference electrodes were placed on the left and right mastoid. Dim light melatonin onset (DLMO) was assessed through 8 hourly saliva samples, from 7 hours before HSon onwards. After the last saliva sample, the FD protocol started with 5 hours for sleep. Participants were woken up in dim or bright polychromatic white light and remained awake under these light conditions for the next 13 hours (Fig 1). Two-hourly test sessions were performed during wakefulness, starting 20 minutes after awakening. Saliva samples to determine melatonin and cortisol concentrations were taken hourly during wakefulness. Iso-caloric snacks were provided immediately after completion of each test session, with caloric value based on estimated basic metabolic rate (BMR) according to the following equation: *BMR* = ((10 ∗ weight(kg)) + (6.25 ∗ *length*(*cm*)) − (5 ∗ *age*(*yrs*)) + 5)) ^20^. After 13 hours of wakefulness under dim or bright light, participants were instructed to go to bed after which the light was switched off. The length of the FD paradigm should cover an integer number of beats of the two involved rhythms. This is needed for the mathematical disentanglement of wake duration related variation from circadian variation. Here, the 18-h FD cycle, consisting of 5 h for sleep and 13 h for wakefulness, was repeated 4 times, resulting in a 72-hour forced desynchrony protocol (4 times 18-h exactly matches 3 times 24-h). After completion, subjects remained in DL in the human isolation facility for an additional sleep opportunity of 3 hours. From 9 hours before the following HSon until 2 hours after that HSon, saliva samples for DLMO determination were collected. After the last sample, subjects were allowed to go home while wearing wrist Actigraphy and returned after at least 3 weeks to participate in the same protocol under opposite light conditions. The order of light conditions was counterbalanced. The experiment was conducted between February and May 2018.

**Figure 1.**
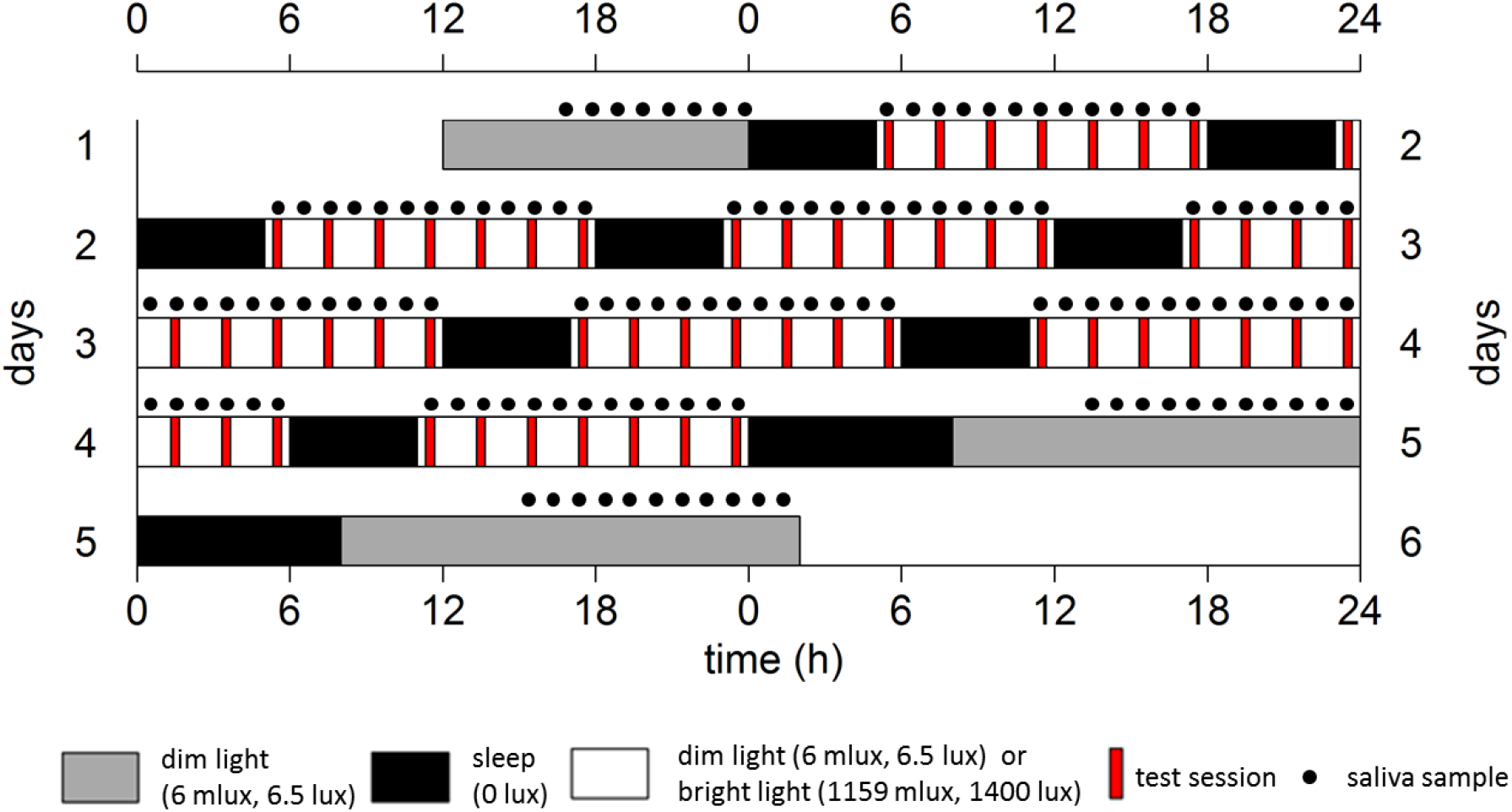
Schematic representation of the experiment design, for an individual with a HSon at 00:00 h, double-plotted. Gray bars indicate dim light (6.5 lux) conditions preceding/following the FD protocol. Black bars represent intervals for sleep (5h, except for the last sleep attempt which is allowed to last for 8 h) with lights off (0 lux), while white bars represent wakefulness in either dim (6 melanopic lux, 6.5 lux) or bright (1159 melanopic lux, 1400 lux) conditions. Black dots indicate saliva samples for melatonin and cortisol assessments, red bars indicate test sessions. After completion of a test session, subject received an iso-caloric snack. The protocol lasted for 72 hours, therefore comprising a full beat (3 × 24h = 72h, 4 × 18h = 72h)

### Test session

Test sessions started with the EEG based indices of alertness ^21^, in which alpha and theta power density during eyes-closed (2 minutes) and open (6 minutes, while focusing on a fixation mark) were determined. Thereafter, subjects completed the Karolinska Sleepiness Scale (KSS; ^21^), which was followed by two auditory reaction time tasks, consisting of a 5-minute Psychomotor Vigilance (PVT) and a 5-minute Go-NoGo (GNG) task, to assess sustained attention ^22^ and executive control ^23^ respectively.

### Light Exposure

Polychromatic white dim light (DL; 6 melanopic lux, 6.5 lux) and bright light (BL; 1159 melanopic lux, 1400 lux) were provided via ceiling-mounted Philips fluorescent light tubes (see extended data Table S1 and Fig S3 for all α-opic illuminance values and spectral composition). Each participant was exposed to the same light intensity throughout the light phase of one FD protocol.

### Data pre-processing: hormone analysis

Saliva was collected using Sarstedt Salivettes with a cotton swab (Salivette, Sarstedt BV, Etten-Leur, the Netherlands). Melatonin concentrations were assessed by radioimmunoassay (RIA) (RK-DSM2; Bühlmann Laboratories AG, Schönenbuch, Switzerland). The limit of detection was 0.5 pg/ml with 9.1% intra-essay variation, and 18.1% inter-essay variation in lowest- and 14.1% in highest concentrations. Cortisol concentrations were detected with RIA analysis (CORT-C2; Cisbio, Bioassays, Parc Marcel Boiteux, Codolet, France), including Bio-rad Immunoassay Plus Control (Lyphochek). The limit of detection was 1 nmol/l, with 10.4% intra-essay and 9.7% inter-essay variation at low- and 3.4% at high cortisol concentrations.

### Data pre-processing: tau estimation

Internal circadian period was estimated by melatonin data, measured as DLMO preceding and following the FD protocol. DLMO was defined as crossing of the 3 pg/ml concentration, by linear interpolating of raw values before and after the cutoff point.

### Data pre-processing: PVT and GNG

A response was defined as anticipation error if it occurred before the time calculated as the average response time of all test sessions minus two standard deviations. Similarly, errors of omission were responses occurring after the time calculated as the average of all test sessions + two standard deviations. These definitions have been shown to be the preferred method over a set cutoff point, since it yields a more accurate representation of individually varying reaction times ^24^. In the GNG-task, errors of commission were characterized as responding to a non-response tone. Other parameters of interest were median RT, average RT, averages of the 10% fastest and 10% slowest RTs, and the average RT in the first and last minute of the 5-minutes test.

### Data pre-processing: EEG based indices of alertness

EEG data were collected using the Temec EEG (Vitaport, 28 channels) system. Electrode impedance was maintained below 5 kΩ. Data was analyzed using Vitascore V1.60. Artefacts were manually removed, after applying high-pass (0.5 Hz) and low-pass (30 Hz) digital filters. Alpha (8.0–12.0 Hz) and theta (4.0–7.9 Hz) power values during eyes-closed (2 min) and eyes-open (5 min) conditions were calculated using Fast Fourier transform (FFT) with a 4 seconds bin width. Power spectra were calculated for every 30 second epoch of EEG data on every derivate.

### Statistics

RStudio (version 1.0.136) was used for statistics and graphics. Wake duration related contributions to neurobehavioral performance measures and subjective alertness assessments were quantified by grouping and averaging data in 2-h bins according to time since sleep offset, starting 0.5h after waking until 12h later. Wake duration related regulation of hormonal parameters such as melatonin and cortisol were quantified on the basis of hourly bins. To determine circadian variation, original data were calculated as a function of circadian phase (in degrees relative to DLMO) in 30-degree bins. Linear mixed models were constructed with light condition as independent variable, time since sleep offset and circadian phase as a fixed effect (categorical variable) and added interaction terms between time since sleep offset and circadian phase, time since sleep offset and light, and circadian phase and light condition. Subject ID and visit were included as a random effects to control for between subject variation and possible order effects. Critical two-sided significance level alpha was 0.05 for all statistical tests. Significant interaction terms (p<0.05) between light- and time effects allowed for calculations of significant light effects by contrast analyses for all combinations of circadian time and time since sleep offset. Significant contrasts were depicted in 3-dimensional graphs, in which circadian variation (in 60 degree bins) is depicted on the x-axis, wake duration related variation on the y-axis, and bright light scores were subtracted from dim light scores, with colors indicating the direction and magnitude of the light effect. Overall effects are indicated in tables. Order of light exposure was non-significant in all models(p<0.05).

## Results

### Melatonin and cortisol

Internal period (*τ*) based on melatonin values increased significantly (on average 21 min) after bright light (BL) exposure (dim light (DL); average ± SEM; 24.25 ± 0.09, BL; average ± SEM; 24.60 ± 0.11, p=2·10^−4^, Fig S4). Melatonin rhythms showed robust oscillations in DL, but disrupted rhythms in BL due to light induced melatonin suppression (Fig 2A, Table 1). There was an effect of time awake on melatonin levels, with significant differences between light conditions (Fig 2B, Table 1). Significant differences between BL and DL also existed depending on circadian clock time, showing dampened melatonin rhythms in BL, but significant variation in DL (Fig 2C, Table 1). These effects were even more apparent in 3D plots, with significantly higher melatonin values after DLMO (phase 0) in the DL condition, without significant differences at circadian phases when melatonin was absent (Fig 2D).

**Table 1:** Summary of statistics of wake duration related variation (process S), circadian variation (process C), interaction between process S and C, and additive effects of bright light exposure. Values from linear mixed models on melatonin and cortisol concentrations, subjective alertness scores, performance (PVT and GNG) and EEG based indices of alertness (Alpha power eyes open and closed, and Theta power with eyes open andclosed).

|  | Wake duration related variation<br>(process S) |  | Circadian variation<br>(process C) |  | Interaction<br>(process S x C) |  | Additive effect of bright<br>light |  |
| --- | --- | --- | --- | --- | --- | --- | --- | --- |
| Melatonin | $F_{(12, 672)}$,<br>$p$ | **7.92,**<br>**<0.00001** | $F_{(5, 672)}$,<br>$p$ | **77.12,**<br>**<0.00001** | $F_{(60, 672)}$,<br>$p$ | **2.14,**<br>**<0.00001** | $F_{(1, 672)}$,<br>$p$ | **152.81,**<br>**<0.00001** |
| Cortisol | $F_{(12, 672)}$,<br>$p$ | **3.37,**<br>**<0.0001** | $F_{(5, 672)}$,<br>$p$ | **75.36,**<br>**<0.00001** | $F_{(60, 672)}$,<br>$p$ | **1.66,**<br>**<0.01** | $F_{(1, 672)}$,<br>$p$ | 0.22,<br>>0.05 |
| Subjective alertness | $F_{(6, 392)}$,<br>$p$ | **17.04,**<br>**<0.0001** | $F_{(5, 392)}$,<br>$p$ | **3.80,**<br>**<0.01** | $F_{(30, 392)}$,<br>$p$ | 0.91,<br>>0.05 | $F_{(1, 392)}$,<br>$p$ | 2.45,<br>>0.05 |
| PVT reaction time | $F_{(6, 392)}$,<br>$p$ | 1.10,<br>>0.05 | $F_{(5, 392)}$,<br>$p$ | 0.00,<br>>0.05 | $F_{(30, 392)}$,<br>$p$ | 0.00,<br>>0.05 | $F_{(1, 392)}$,<br>$p$ | **113.69,**<br>**<0.00001** |
| GNG reaction time | $F_{(6, 392)}$,<br>$p$ | 1.76,<br>>0.05 | $F_{(5, 392)}$,<br>$p$ | 0.00,<br>>0.05 | $F_{(30, 392)}$,<br>$p$ | 0.00,<br>>0.05 | $F_{(1, 392)}$,<br>$p$ | **100.19,**<br>**<0.00001** |
| Alpha power eyes open | $F_{(6, 392)}$,<br>$p$ | 0.77,<br>>0.05 | $F_{(5, 392)}$,<br>$p$ | 1.44,<br>>0.05 | $F_{(30, 392)}$,<br>$p$ | 0.64,<br>>0.05 | $F_{(1, 392)}$,<br>$p$ | **20.69,**<br>**<0.00001** |
| Alpha power eyes closed | $F_{(6, 392)}$,<br>$p$ | 0.56,<br>>0.05 | $F_{(5, 392)}$,<br>$p$ | 1.11,<br>>0.05 | $F_{(30, 392)}$,<br>$p$ | 0.44,<br>>0.05 | $F_{(1, 392)}$,<br>$p$ | **34.24,**<br>**<0.0001** |
| Theta power eyes open | $F_{(6, 392)}$,<br>$p$ | 0.49,<br>>0.05 | $F_{(5, 392)}$,<br>$p$ | 1.20,<br>>0.05 | $F_{(30, 392)}$,<br>$p$ | 0.41,<br>>0.05 | $F_{(1, 392)}$,<br>$p$ | 0.01,<br>>0.05 |
| Theta power eyes closed | $F_{(6, 392)}$,<br>$p$ | 1.48,<br>>0.05 | $F_{(5, 392)}$,<br>$p$ | 0.92,<br>>0.05 | $F_{(30, 392)}$,<br>$p$ | 0.89,<br>>0.05 | $F_{(1, 392)}$,<br>$p$ | 0.30,<br>>0.05 |

**Figure 2:**
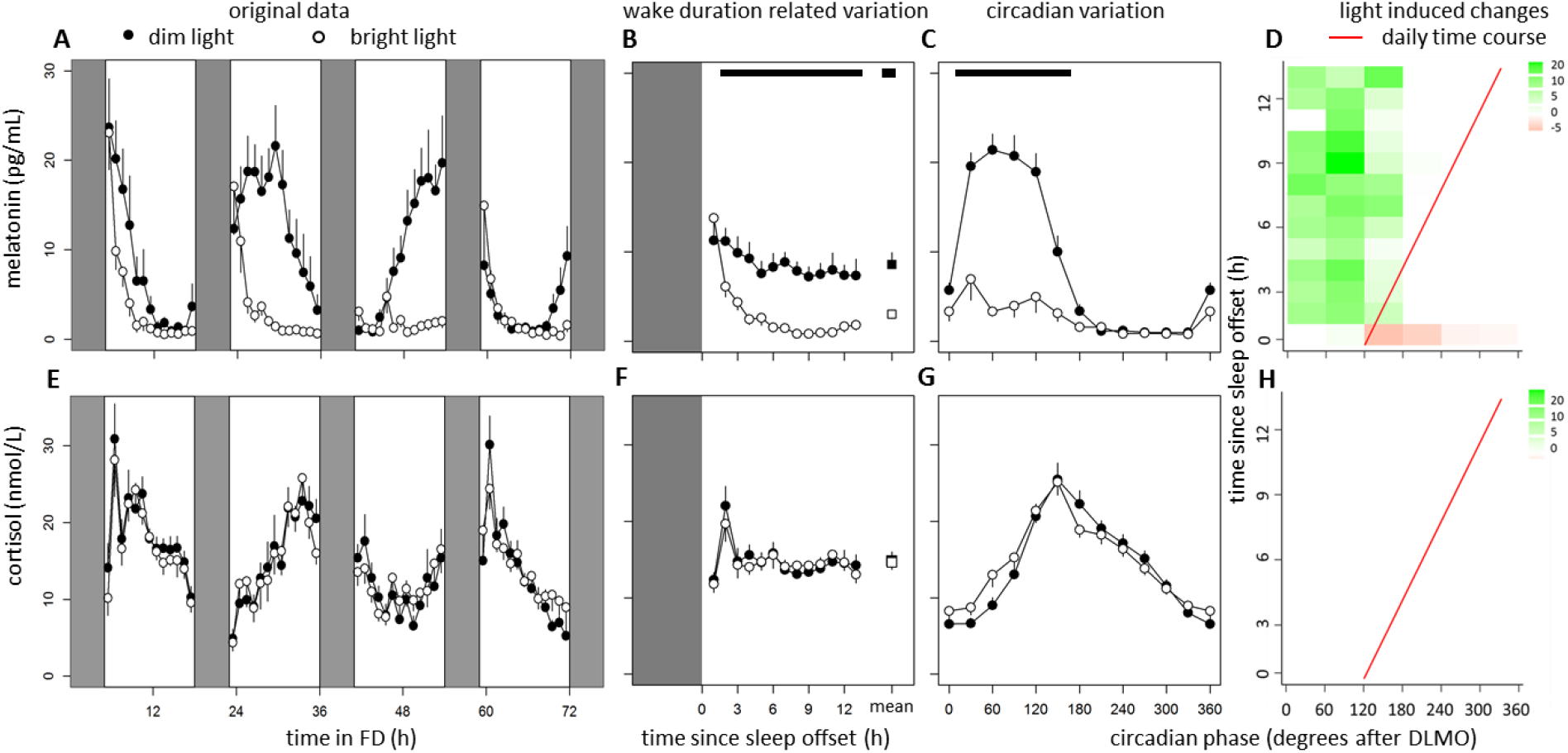
Data of melatonin (top panel) and cortisol (bottom panel) concentrations. Time course of melatonin (A) and cortisol (E) during the FD protocol. Data replotted as time since sleep offset (B and F) and circadian phase in degrees after DLMO (C and G), for melatonin and cortisol respectively. Contrast analysis describing light induced decrease for all combinations of circadian clock phase and time since sleep offset for melatonin (D) and cortisol (H). Data represent mean ± standard error of the mean, with 7 subjects per group. Black dots indicate data collected in dim light, white dots represent data collected in bright light and black and white squares represent averages over all data points under DL and BL respectively. Red line indicates the expected time course over a regular day. Shaded areas represent scheduled sleep (at 0 lux). Significant differences between light conditions (p<0.05) are indicated by horizontal black bars (B,C, F, G) or colored rectangles (D, H).

Successive cortisol cycles indicate free running of the circadian clock, without significant differences between light conditions (Fig 2E, Table 1). There were significant effects of time awake (Fig 2F, Table 1). Cortisol levels varied significantly with circadian clock time, independent of light condition (Fig 2G, Table 1). There were no significant interaction effects between time and light condition (Fig 2H).

### Relative coordination

To investigate possible relative coordination, residual cortisol data was further analyzed after subtraction of both the calculated wake duration related- and circadian variation. Results indicated that these data did not demonstrate significant relative coordination, since a fitted 72-h sine wave was not significant (DL: F_2,359_=0.14, p=0.87 and BL: F_2,359_=0.17, p=0.84; Fig. S5). The residual variation was respectively 1.61 and 1.83% of the variation in the raw data under DL and BL conditions, indicating that the calculated homeostatic and circadian clock time fluctuations explain almost all of the variation in cortisol concentration.

### Subjective alertness

Subjective alertness scores (Fig 3A) increased with time awake, and the onset of subjective sleepiness was significantly delayed by BL exposure (Fig 3B). Circadian variation was established, independent of light exposure (Fig 3C, Table 1). Significant interactions were found between circadian phase, time since sleep offset, and light induced change in subjective alertness, predominantly after DLMO, but not during the daily time course (Fig 3D).

**Figure 3:**
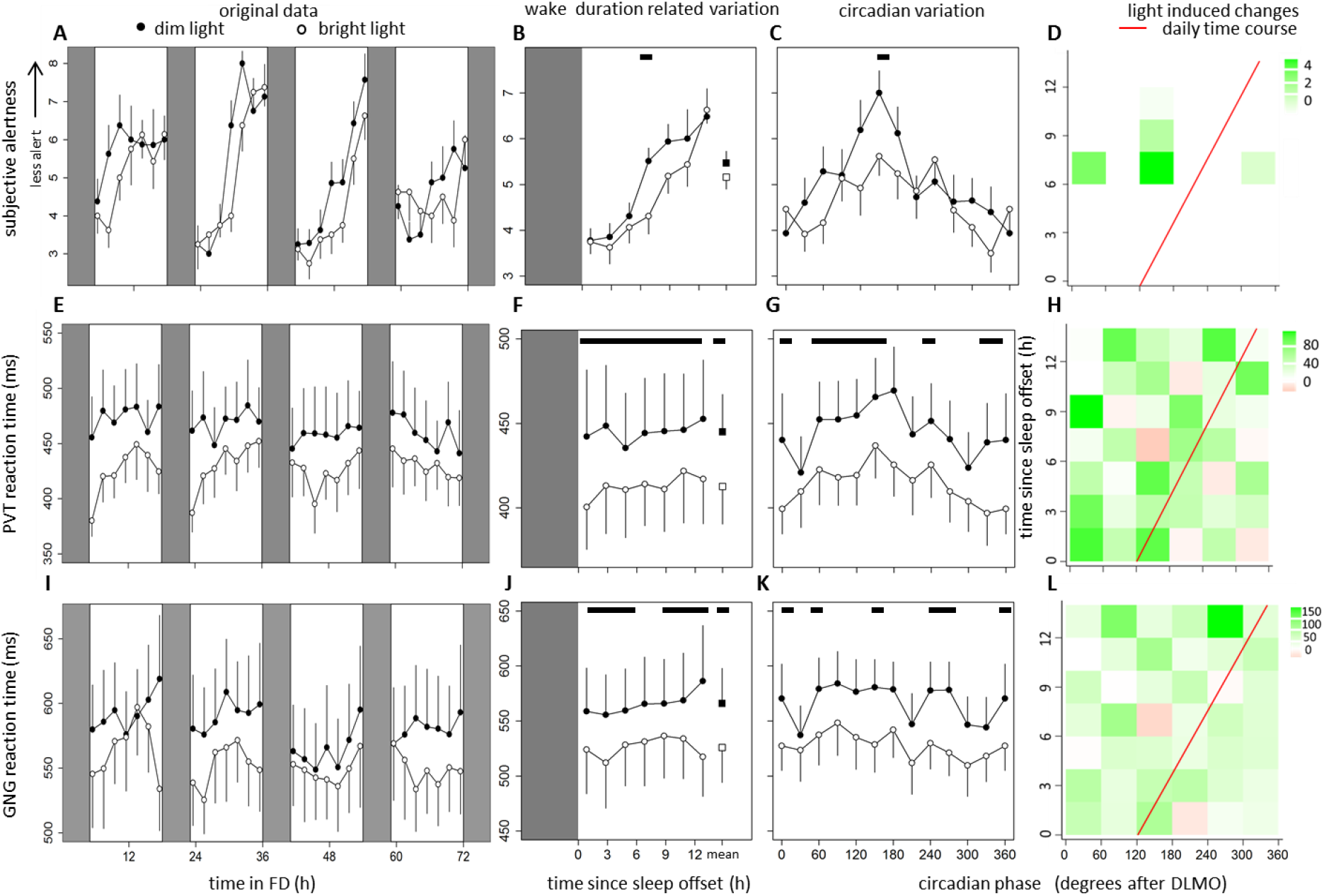
Data of subjective alertness (top panel), average reaction time on the PVT (middle panel) and GNG (bottom panel). Time course of subjective alertness (A), performance on the PVT (E) and GNG (I) during the FD protocol. Data replotted as time since sleep offset (B, F and J) and circadian phase in degrees after DLMO (C, G and K), for subjective alertness, PVT performance and GNG performance respectively. Contrast analysis describing light induced decrease for all combinations of circadian clock phase and time since sleep offset for subjective sleepiness (D), PVT performance (H) and GNG performance (L). Data represent mean ± standard error of the mean, with 7 subjects per group. Black dots indicate data collected in dim light, white dots represent data collected in bright light and black and white squares represent averages over all data points under DL and BL respectively. Red line indicates the expected time course over a regular day. Shaded areas represent scheduled sleep (at 0 lux). Significant differences between light conditions (p<0.05) are indicated by horizontal black bars (B,C, F, G, J, K) or colored rectangles (D, H, L).

### Performance: PVT

Sustained attention (defined as average reaction time on the PVT (ms), Fig 3E) was better in BL compared to DL, independent of time awake (Fig 3F, Table 1). There was no significant effect of circadian clock time, but there were significant differences between light conditions (Fig 3G, Table 1). Significant interactions were found between circadian phase, time since sleep offset, and light induced change in performance, with significant light induced improvements in reaction time occurring during the daily time course (Fig 3H). Similar effects were found in detailed parameters of the PVT task (Fig S6, Table S2).

### Performance: Go-NoGo

Executive performance (assessed with the Go-NoGo task, Fig 3I), was better in BL independent of time awake (Fig 3J, Table 1). There was no significant effect of circadian clock time, which was affected by light exposure (Fig 3K, Table 1). There were significant interactions between circadian phase, time since sleep offset, and light induced change in performance (Fig 3L). Similar effects were found in detailed parameters of this task (Fig S7, Table S2).

### EEG based indices of alertness

Wake EEG analysis was measured in the electrode placed at central location (Fig 4A,E,I,M). Data revealed a significant effect of light exposure on alpha power with eyes open independent of time awake (Fig 4B, Table 1). Circadian clock time did not significantly affect EEG power density, although this was significantly decreased by BL exposure (Fig 4C, Table 1). Significant interactions were detected between circadian phase, time since sleep offset, and light induced change in EEG based indices of alertness, some with significant lower alpha power occurring during the daily time course following bright light exposure (Fig 4D). EEG activity in the alpha band when eyes were closed was independent of time awake, but significantly decreased by BL exposure (Fig 4F, Table 1). There were no significant effects of circadian clock time, but significant decreases due to BL exposure were found (Fig 4G, Table 1). There were no significant interactions between circadian phase, time since sleep offset, and light induced changes in alpha power with eyes closed during the daily time course (Fig 4H). When eyes were open, theta activity was not significantly affected by time awake or light exposure (Fig 4J, Table 1). There were no significant effects of circadian clock time or light exposure (Fig 4K, Table 1), and no significant interactions between circadian phase, time since sleep offset, and light induced changes in theta power with eyes open during the daily time course (Fig 4L). When eyes were closed, theta band activity was not significantly affected by time awake or light exposure (Fig 4N, Table 1). There were no significant effects of circadian clock time, or light intervention (Fig 4O), nor of a significant interaction between circadian phase, time since sleep offset, and light induced changes in theta power with eyes closed during the daily time course (Fig 4P). Data from other electrode positions can be found in S8, S9 and Table S3. Correlations between the various measures of alertness can be found in Fig S10. All results are summarized in Table 2.

**Table 2:** Summary of effects of wake duration related variation, internal clock time, the interaction between both processes, direction of additive effects of bright light exposure, and interaction effects between light exposure and timing of exposure during the daily time course. ‘Y’ and ‘N’ indicate significant or non-significant effects respectively, ‘+’ indicates a significant increase, ‘−‘ represents a significant decrease, and non-significant changes are indicated by ‘0’.

|  | Wake duration related variation<br>(process S) | Internal clock time<br>(process C) | Interaction<br>(process S x C) | Additive effect<br>of bright light | Interactive effect during<br>daily time course |
| --- | --- | --- | --- | --- | --- |
| Melatonin | Y | Y | Y | - | N |
| Cortisol | Y | Y | Y | 0 | N |
| Subjective alertness | Y | Y | N | 0 | N |
| PVT reaction time | N | N | N | - | Y |
| GNG reaction time | N | N | N | - | Y |
| Alpha power eyes open | N | N | N | - | Y |
| Alpha power eyes closed | N | N | N | - | N |
| Theta power eyes open | N | N | N | 0 | N |
| Theta power eyes closed | N | N | N | 0 | N |

**Figure 4:**
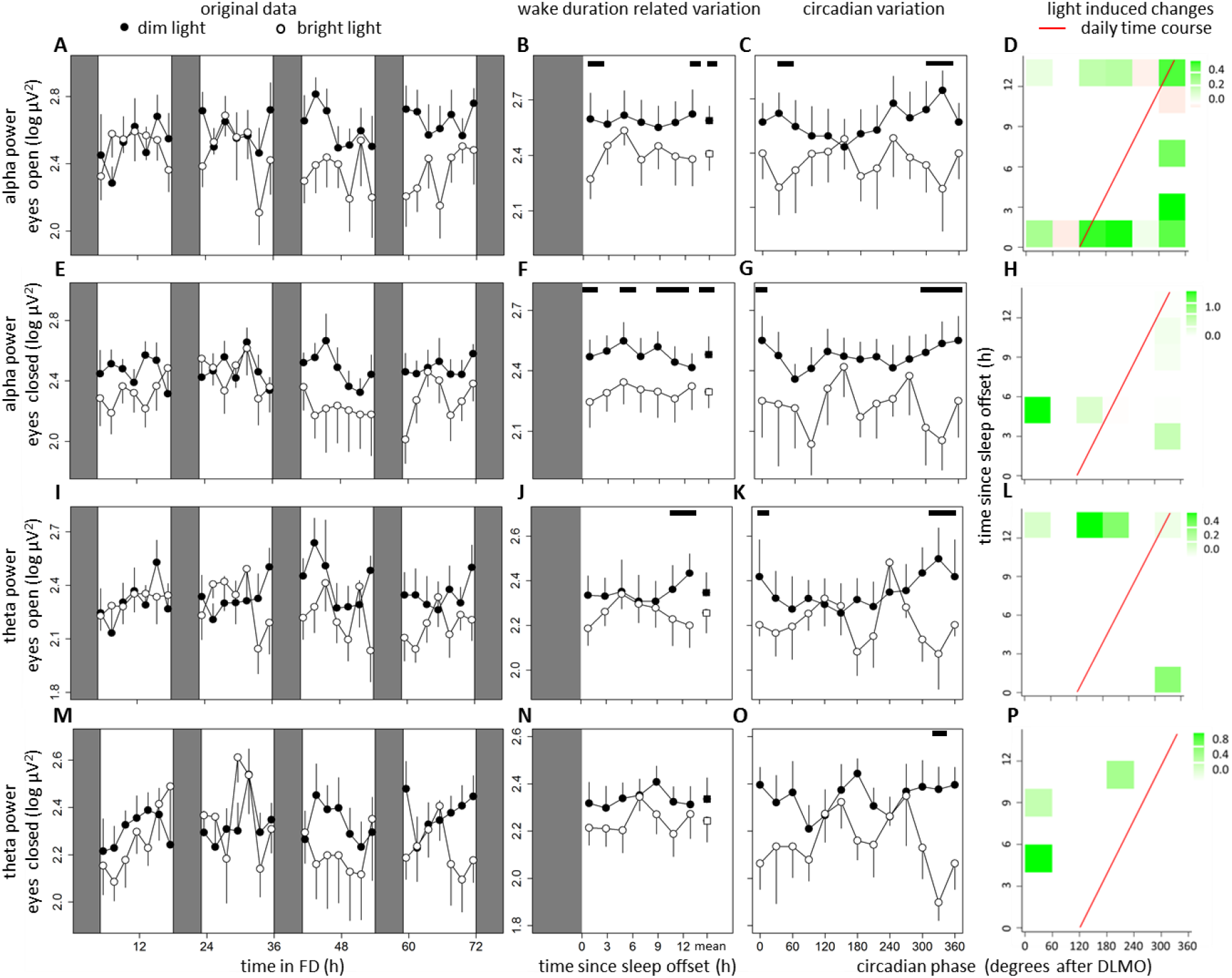
Data of alpha power with eyes open and closed (top panels) and theta power with eyes open and closed (bottom panels). Time course of alpha power with eyes open (A), and closed (E), as well as theta power with eyes open (I) and closed (M) during the FD protocol. Data replotted as time since sleep offset (B, F, J, and N) and circadian phase in degrees after DLMO (C, G, K and O), for alpha power with eyes open and closed, as well as theta power with eyes open and closed respectively. Contrast analysis describing light induced decrease for all combinations of circadian clock phase and time since sleep offset for alpha power with eyes open (D), and closed (H) as well as theta power with eyes open (L), and closed (P). Data represent mean ± standard error of the mean, with 7 subjects per group. Black dots indicate data collected in dim light, white dots represent data collected in bright light and black and white squares represent averages over all data points under DL and BL respectively. Red line indicates the expected time course over a regular day. Shaded areas represent scheduled sleep (at 0 lux). Significant differences between light conditions (p<0.05) are indicated by horizontal black bars (B,C, F, G, J, K, N, O) or colored rectangles (D, H, L,P).

## Discussion

Light can increase (subjective) alertness during the night, but results are less conclusive during daytime ^9^. Both wake duration related variation and circadian clock phase might contribute to this discrepancy. In the current FD study, we found that light can increase objective measures of alertness independent of time awake, while positive effects of light on subjective alertness is only reported 6-9 hours after waking up (Fig.3). High intensity light exposure effectively postponed the increase of subjective alertness with progressing time awake, although light induced improvements could not be determined during the daily time course (Fig.3). Performance was better throughout the wake interval in high intensity light, independent of sleep pressure build-up or circadian clock phase, with significant improvements during the daily time course. There were no significant effects of time awake or circadian variation on EEG based indices of alertness, although there were light induced decreases. Overall, these did not occur during the daily time course.

### Bright light FD design and cortisol as a phase marker

Dim light (<10 lux) has been standard procedure since the first FD experiment was performed in ^13^. Apart from model calculations ^15^, this is the first study we know that shows that short FD protocols can also be used for disentangling effects of bright light gated by wake duration related variation and circadian processes. During FD paradigms, concentrations of the nocturnal hormone melatonin have regularly been used as output measures of SCN rhythmicity. Cortisol had to be used as a circadian phase marker during the experiment instead of the usual usage of melatonin because high intensity light suppresses melatonin production ^25^. A free-running 24-hour cortisol rhythm with uniform phase progression in both light conditions suggests that cortisol can be used as a reliable output measure of the circadian clock under these conditions, and that there was no indication of relative coordination under high light intensity exposure. Suggestions that light might decrease ^26–28^ or increase ^29^ cortisol in humans are not confirmed by our data and others^30–32^. The absence of the cortisol awakening response in combinations of circadian clock time and wake duration related variation employed in the current FD protocol may have contributed to the absence of light effects on cortisol.

### Lengthening of the internal period

Based on the dim light melatonin onset preceding and following the FD protocol, the internal circadian period showed a 21-min increase under BL (Fig. S4). This is in line with lengthening of circadian period with increasing continuous light intensity as observed in most mammals, including diurnal primates ^33,34^. On the other hand, diurnal vertebrates overall seem to decrease circadian period under the influence of light, as was described by Aschoff’s rule and his own findings in humans ^35,36^. Earlier model simulations indicated that FD protocols with limited amount of cycles may typically overestimate circadian period at high light intensities ^15^, and should therefore not be over-interpreted.

### Subjective alerting effects of light depend on homeostatic control

After the circadian signal is subtracted from the original data, our data confirm subjective alertness increases with time awake ^12^. Subjective alertness ratings were similar under both light conditions, both at the start and the end of the light phase, suggesting that the decrease in alertness is postponed by high intensity light exposure. Subjective alertness scores after awakening were similar to natural conditions ^37^, indicating that 5 h of sleep, 13 h wakefulness did not increase sleep pressure. Moreover, the relationship between alertness scores and alerting effects of light based on homeostatic sleep pressure suggest a parabolic function; a ceiling effect is present after awakening, and as a consequence alertness cannot improve beyond its limits, but with increasing sleep pressure build-up, alertness decreases, creating room for improvement by light. When sleep pressure rises further, accumulated sleep pressure overrides positive effects of bright light exposure. This hypothesis is consistent with the observations that alerting effects of light during daytime are hard to determine in well-rested individuals ^38^, while significant effects have been reported in sleep deprived individuals ^39^. In addition to effects of time awake, there is also circadian variation in subjective alertness, with the largest decrease in alertness at the end of the night, as has been reported in other FD studies ^12^. Most likely, this contributes to sleep maintenance when most sleep pressure has dissipated ^12,40^. Likewise, the circadian contribution to alertness at the end of the day compensates for the increased homeostatic sleep pressure at that time. Noteworthy is that although light has significant effects on subjective alertness, none of these occur during the daily time course (Fig 3D).

### Objective alerting effects of light are independent of homeostatic control

Performance was significantly improved under bright light, but did not change as a function of time of day. This is in contrast with some ^11,41^ but not all literature ^42,43^, in which significant effects of time awake have been described. Similar patterns have been reported in EEG based indices of alertness ^44^. Although significant effects of bright light exposure in alpha activity were determined, wake duration- or circadian clock phase related effects could not be assessed. A possible explanation for discrepancies between our and other studies might be varying lengths of FD protocols. Longer FD cycles induce higher sleep pressure levels and therefore larger declines in performance and increases in alpha-theta activity. Moreover, accumulating sleep pressure may differently affect subjective alertness, performance and EEG ^45^, as these measures do not always correlate well together ^46,47^. Our data adds to the body of literature supporting this, since both performance and wake EEG reflect similar patterns, which do not coincide with subjective alertness patterns. It is noteworthy that significant improvements in performance were present after merely 20 minutes of high intensity light exposure on the first day of the FD paradigm, suggesting that light effects on these parameters are relatively fast. Light exposure may counteract initial decrements in performance due to sleep inertia after awakening.

### Variation over the daily time course

Data collected here indicate that over the time course of a normal daily wake period, light can improve objective, but not subjective parameters of alertness (Fig 3). This data suggest that bright light exposure at daytime is particularly beneficial for performance, although this benefit may not be subjectively perceived.

In conclusion, this is the first study to investigate effects of light on alertness under forced desynchrony conditions, indicating that (1) our short forced desynchrony protocol can be used for disentangling wake duration related variation and circadian aspects under high intensity lighting, (2) bright light can postpone the onset of subjective alertness, depending on wake duration induced variation, and (3) performance and is improved by high intensity light independent of wake duration related variation or circadian clock phase. This suggests that light during office hours might be beneficial for performance, even though this may not be perceived as such.

## Acknowledgements

We thank all subjects for participation. We also thank Bonnie de Vries for assistance with melatonin and cortisol radioimmunoassay analysis. This research was funded by the University of Groningen Campus Fryslân (Grant No. 01110939; co-financed by Philips Drachten and Provincie Fryslân).

## Author contribution

R.L., T.W., M.v.K., K.O., S.G.F. were involved in data acquisition. R.L. and T.W. conducted data analysis. R.L. drafted the manuscript. Authors R.L., T.W., D.G.M.B., R.A.H. contributed to the concept design, interpretation of data and editing of the original manuscript.

## Conflict of interest

The authors do not declare any conflict of interest.

## Notes

### Competing Interest Statement

The authors have declared no competing interest.

## References

1. Moore, R. Y. & Eichler, V. B. Loss of a circadian adrenal corticosterone rhythm following suprachiasmatic lesions in the rat. Brain Res. Bull. 72, 54–56 (1972).

2. Stephan, F. K. & Zucker, I. Circadian rhythms in drinking behavior and locomotor activity of rats are eliminated by hypothalamic lesions. Proc. Natl. Acad. Sci. USA 69, 1583–1586 (1972).

3. Cajochen, C., Zeitzer, J. M., Czeisler, C. A. & Dijk, D. J. Dose-response relationship for light intensity and ocular and electroencephalographic correlates of human alertness. Behav. Brain Res. 115, 75–83 (2000).

4. Posner, M. I. Measuring alertness. Ann. N. Y. Acad. Sci. 1129, 193–199 (2008).

5. Figueiro, M. G., Sahin, L., Wood, B. M. & Plitnick, B. A. Light at Night and Measures of Alertness and Performance: Implications for Shift Workers. Biol. Res. Nurs. 18,(2015).

6. Curcio, G., Casagrande, M. & Bertini, M. Sleepiness: evaluating and quantifying methods. Int. J. Psychophysiol. 41, 251–263 (2001).

7. Alexandre, C. et al. Decreased alertness due to sleep loss increases pain sensitivity in mice. Nat. Med. 23, 768–774 (2017).

8. Pardi, D., Buman, M., Black, J., Lammers, G. J. & Zeitzer, J. M. Eating Decisions Based on Alertness Levels after a Single Night of Sleep Manipulation: A Randomized Clinical Trial. Sleep 40, 1–8 (2016).

9. Lok, R., Smolders, K. C. H. J., Beersma, D. G. M. & de Kort, Y. A. W. Light, Alertness, and Alerting Effects of White Light: A Literature Overview. J. Biol. Rhythms 33,(2018).

10. Aston-Jones, G. Brain structures and receptors involved in alertness. Sleep Med. 6, 3–7 (2005).

11. Wright, K. P., Hull, J. T. & Czeisler, C. A. Relationship between alertness, performance, and body temperature in humans. Am. J. Physiol. - Regul. Integr. Comp. Physiol. 283, 1370–1377 (2002).

12. Dijk, D. J., Duffy, J. F. & Czeisler, C. A. Circadian and sleep/wake dependent aspects of subjective a. alertness and cognitive performance. J. Sleep Res. 1, 112–117 (1992).

13. Kleitman, N. & Kleitman, E. Effect of Non-Twenty-Four-Hour Routines of Living on Oral Temperature and Heart Rate. Am. Physiol. Soc. 6,(1953).

14. Hiddinga, A. E., Beersma, D. G. M. & Van den Hoofdakker, R. H. Endogenous and exogenous components in the circadian variation of core body temperature in humans. J. Sleep Res. 6, 156–63 (1997).

15. Woelders, T., Beersma, D. G. M., Gordijn, M. C. M., Hut, R. A. & Wams, E. J. Daily Light Exposure Patterns Reveal Phase and Period of the Human Circadian Clock. J. Biol. Rhythms 32, 174–286 (2017).

16. Daan, S., Beersma, D. G. M. & Borbély, A. A. Timing of human sleep: recovery process gated by a circadian pacemaker. Am. J. Physiol. 246,R161–R183 (1984).

17. Roenneberg, T., Wirz-Justice, A. & Merrow, M. Life between clocks: Daily temporal patterns of human chronotypes. J. Biol. Rhythms 18, 80–90 (2003).

18. Buysse, D. J. et al. The Pittsburgh Sleep Quality Index: a new instrument for psychiatric practice and research. Psychiatry Res. 28, 193–213 (1989).

19. Ishihara, S. The Series of Plates Designed as a Tests for Colour-Blindness. Nature (1972).

20. Mifflin, M. D. et al. A new predictive equation in healthy individuals3 for resting energy. Am. J. Clin. Nutr. 51, 241–247 (1990).

21. Åkerstedt, T. & Gillberg, M. Subjective and Objective Sleepiness in the Active Individual. Int. J. Neurosci. 52, 29–37 (1990).

22. Dinges, D. F. & Powell, J. W. Microcomputer analyses of performance on a sustained operations. Behav. Res. Methods, Instruments Comput. 17, 652–655 (1985).

23. Barry, R. J., De Blasio, F. M., De Pascalis, V. & Karamacoska, D. Preferred EEG brain states at stimulus onset in a fixed interstimulus interval equiprobable auditory Go/NoGo task: A definitive study. Int. J. Psychophysiol. 94, 42–58 (2014).

24. Gabel, V. Auditory psychomotor vigilance testing in older and young adults : a revised threshold setting procedure. 1021–1025 (2019).

25. Rollag, M. D., Panke, E. S., Trakulrungski, W. Trakulrungski, C. & Reiter, R. J. Quantification of daily melatonin synthesis in the hamsterpineal gland. Encorinology 106,231 (1980).

26. Kostoglou-Athanassiou, I., Treacher, D. F., Wheeler, M. J. & Forsling, M. L. Bright light exposure and pituitary hormone secretion. Clin. Endocrinol. (Oxf). 48, 73–79 (1998).

27. BECK-Friis, J., Borg, G. & Wetterberg, L. Rebound Increase of Nocturnal Serum Melatonin Levels following Evening Suppression by Bright Light Exposure in Healthy Men: Relation to Cortisol Levels and Morning Exposure. Ann. N. Y. Acad. Sci. 453, 371–375 (1985).

28. Jung, C. M. et al. Acute effects of bright light exposure on cortisol levels. J. Biol. Rhythms 25, 208–216 (2010).

29. Scheer, F. A. J. L., van Doornen, L. J. P. & Buijs, R. M. Light and Diurnal Cycle Affect Human Heart Rate: Possible Role for the Circadian Pacemaker. J. Biol. Rhythms 14, 202–212 (1999).

30. Ruger, M., Gordijn, M. C., Beersma, D. G., de Vries, B. & Daan., S. Time-of-day-dependent effects of bright light exposure on human psychophysiology: comparison of daytime and nighttime exposure. AJP Regul. Integr. Comp. Physiol. 290,R1413–R1420 (2005).

31. Mcintyre, I. M., Norman, T. R., Burrows, G. D. & Armstrong, S. M. Melatonin, cortisol and prolactin response to acute nocturnal light exposure in healthy volunteers. 17, 243–248 (1992).

32. Lavoie, S., Paquet, J., Selmaoui, B., Rufiange, M. & Dumont, M. Vigilance levels during and after bright light exposure in the first half of the night. Chronobiol. Int. 20, 1019–1038 (2003).

33. Klerman, E. B., Dijk, D.-J., Kronauer, R. E. & Czeisler, C. A. Simulations pacemaker : of light effects on the human circadian implications for assessment of intrinsic period. Am. J. Physiol. 270, 271–282 (1996).

34. Weber, F. Die Periodenlänge der circadianen Laufperiodizität. Naturwissenschaften 54, 122 (1967).

35. Aschoff, J. Tierische Periodik unter dem Einfluss van Zeitgebern. Z. Tierpsychologie 15, 1–30 (1958).

36. Aschoff, J. Circadian rhythms in man. Science (80-.). 148, 1427–1432 (1965).

37. Åkerstedt, T., Hallvig, D. & Kecklund, G. Normative data on the diurnal pattern of the Karolinska Sleepiness Scale ratings and its relation to age, sex, work, stress, sleep quality and sickness absence/illness in a large sample of daytime workers. J. Sleep Res. 26, 559–566 (2017).

38. Lok, R., Woelders, T., Gordijn, M. C. M., Hut, R. A. & Beersma, D. G. M. White Light During Daytime Does Not Improve Alertness in Well-rested Individuals. J. Biol. Rhythms 33, 637–648 (2018).

39. Phipps-Nelson, J., Redman, J. R., Dijk, D. J. & Rajaratnam, S. M. W. Daytime exposure to bright Light, as compared to dim light, decreases sleepiness and improves psychomotor vigilance performance. Sleep 26, 695–700 (2003).

40. Dijk, D. J. & Czeisler, C. A. Paradoxical timing of the circadian rhythm of sleep propensity serves to consolidate sleep and wakefulness in humans Time of Day. Neurosci. Lett. 166, 63–68 (1994).

41. Silva, E. J., Wang, W., Ronda, J. M., Wyatt, J. K. & Duffy, J. F. Circadian and wake-dependent influences on subjective sleepiness, cognitive throughput, and reaction time performance in older and young adults. Sleep 33, 481–490 (2010).

42. Lee, J. H. et al. Neurobehavioral performance in young adults living on a 28-h day for 6 weeks. Sleep 32, 905–13 (2009).

43. Zhou, X. et al. Dynamics of neurobehavioral performance variability under forced desynchrony: evidence of state instability. Sleep 34, 57–63 (2011).

44. Cajochen, C., Wyatt, J. K., Czeisler, C. A. & Dijk, D. J. Separation of circadian and wake duration-dependent modulation of EEG activation during wakefulness. Neuroscience 114, 1047–1060 (2002).

45. Postnova, S., Lockley, S. W. & Robinson, P. A. Prediction of Cognitive Performance and Subjective Sleepiness Using a Model of Arousal Dynamics. J. Biol. Rhythms 33, 203–218 (2018).

46. Van Dongen, H. P. A., Maislin, G., Mullington, J. M. & Dinges, D. F. The cumulative cost of additional wakefulness: dose-response effects on neurobehavioral functions and sleep physiology from chronic sleep restriction and total sleep deprivation. Sleep 26, 117–126 (2003).

47. Zhou, X. et al. Mismatch between subjective alertness and objective performance under sleep restriction is greatest during the biological night. J. Sleep Res. 21, 40–49 (2012).

