## Supporting Information for "Light effects on circadian and homeostatic regulation: alertness increases independent of time awake"

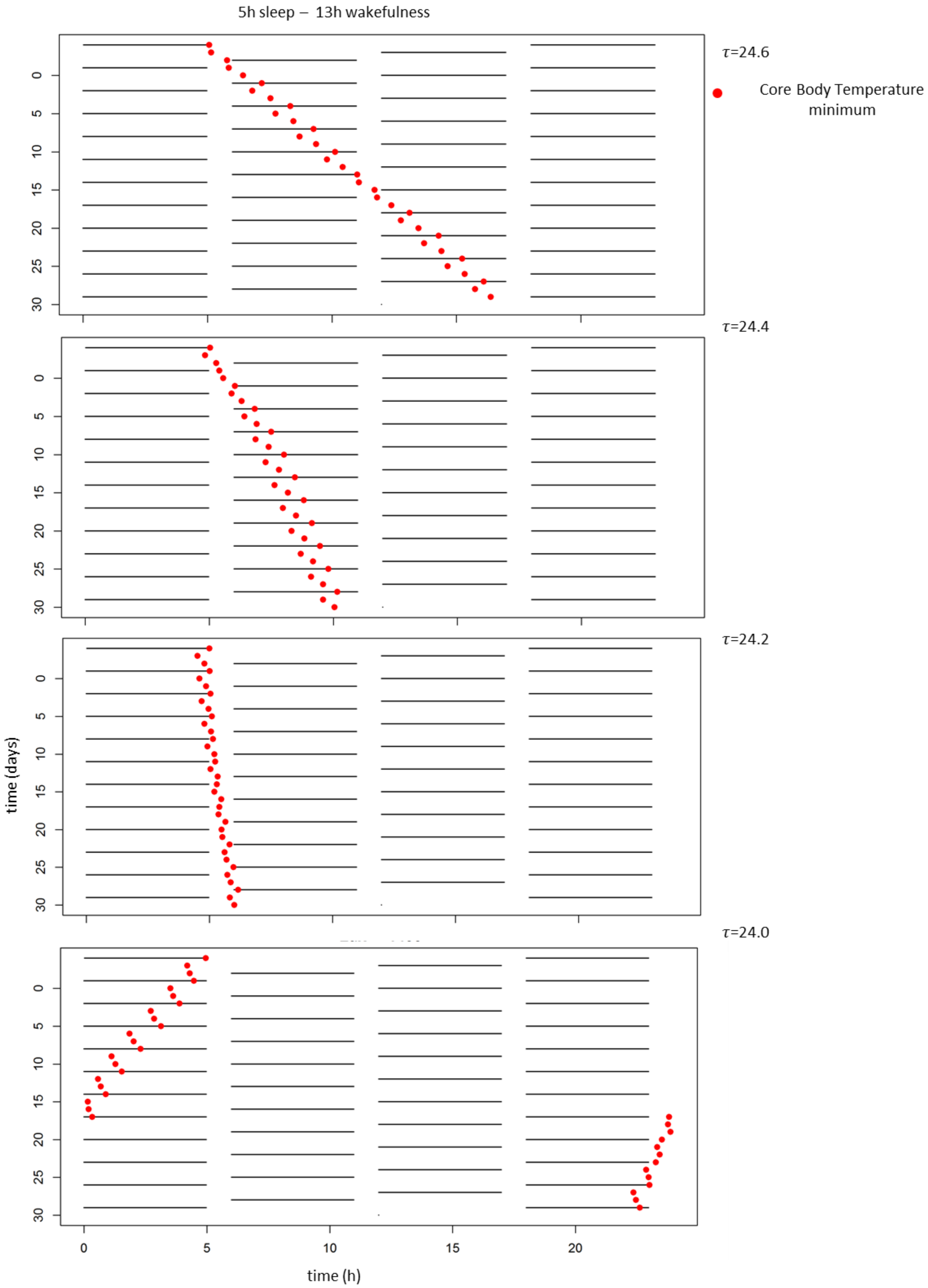


**Figure S1. Simulations of Core Body Temperature minimum, using an adjusted version of Kronauer model under 5h sleep - 13h wakefulness conditions.** Simulations indicate that under 5h sleep and 13h wakefulness, progression of Core Body Temperature minimum as phase marker of the clock is relatively uniform, with a protocol duration of three 24h days. Simulations for three different internal periods ($\tau$)are shown.

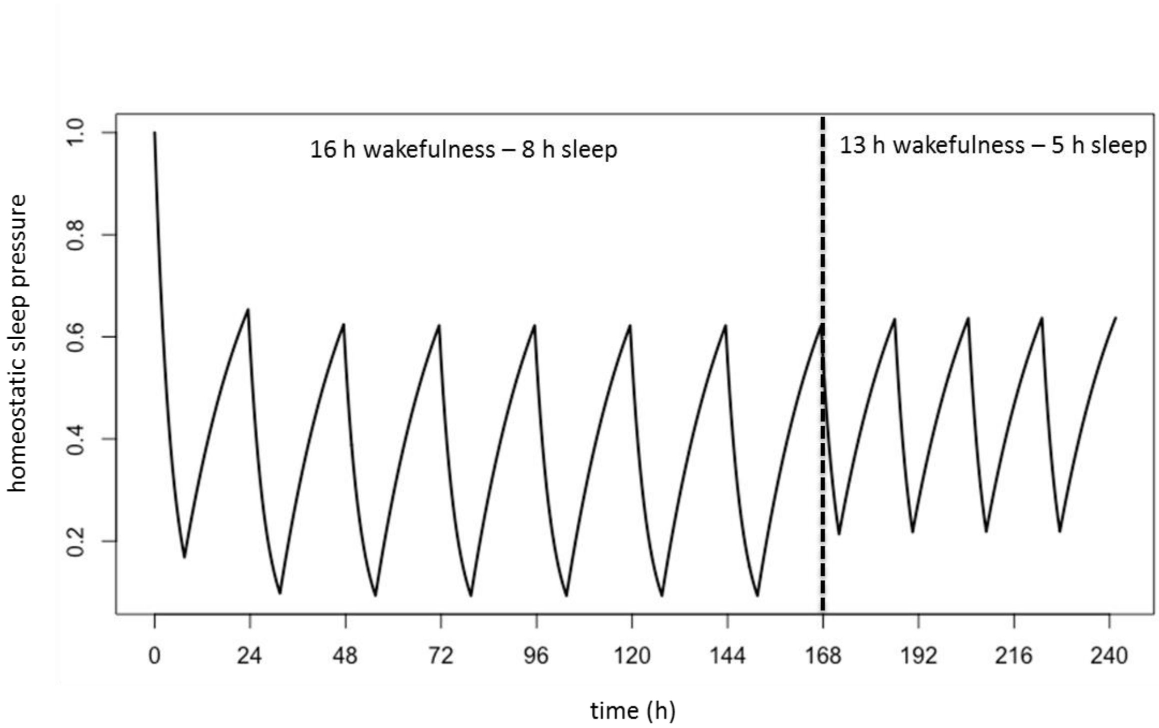


**Figure S2. Simulations of sleep pressure build-up under normal, rested conditions (16h wakefulness – 8h sleep) versus sleep pressure build-up during 13h wakefulness – 5h sleep conditions.** Simulations indicate that under 13h wakefulness and 5h sleep, systematic fluctuations of sleep pressure over the sleep-wake schedule does not change over the course of the FD protocol. Compared to normal sleep timing, sleep pressure in forced desynchrony at awakening is relatively high, while it is about normal at the end of the wake interval.

**Table S1: Photometric properties of dim- and bright light according to the Lucas file(Lucas et al., 2014)**

| Type | Peak irradiance (nm) | Illuminance (lux) | Cyanopic (lux) | Melanopic (lux) | Rhodopic (lux) | Chloropic (lux) | Erythropic (lux) |
|---|---|---|---|---|---|---|---|
| Dim light | 545 | 6.5 | 7.01 | 5.88 | 6.03 | 6.28 | 6.50 |
| Bright light | 545 | 1400 | 1433.41 | 1159.27 | 1189.46 | 1247.15 | 1297.23 |

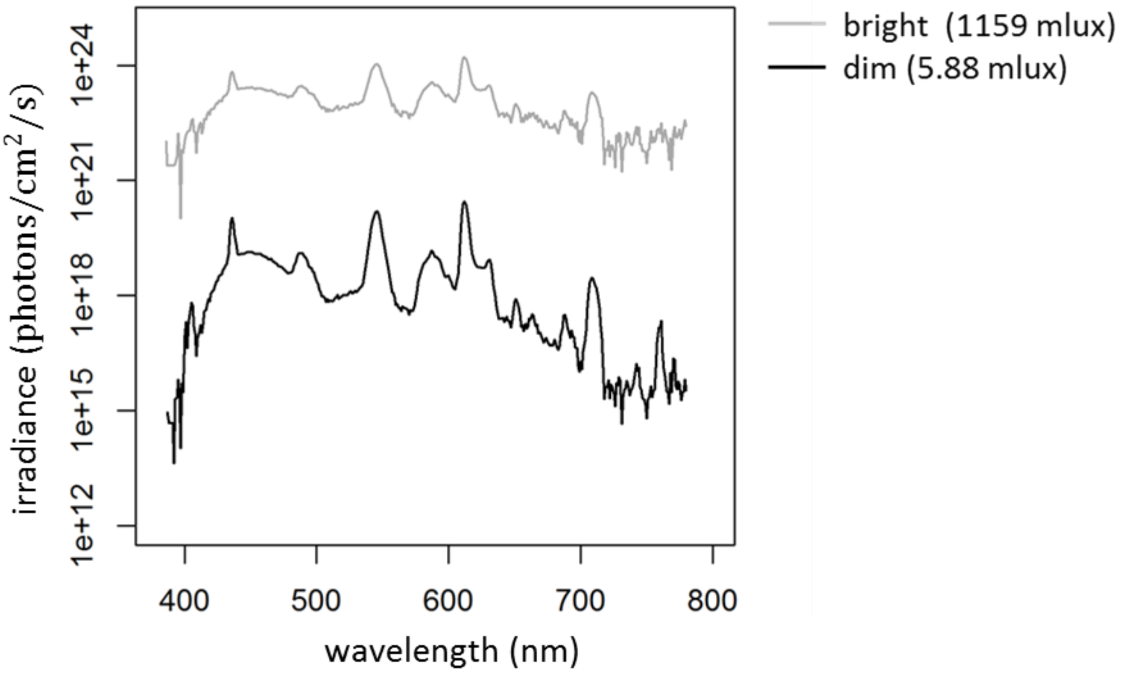


**Figure S3: Spectral composition of dim- (black) and bright light (grey).** Illuminance was measured on the vertical plane at the level of the eye. Light was generated with ceiling-mounted Philips fluorescent light tubes.

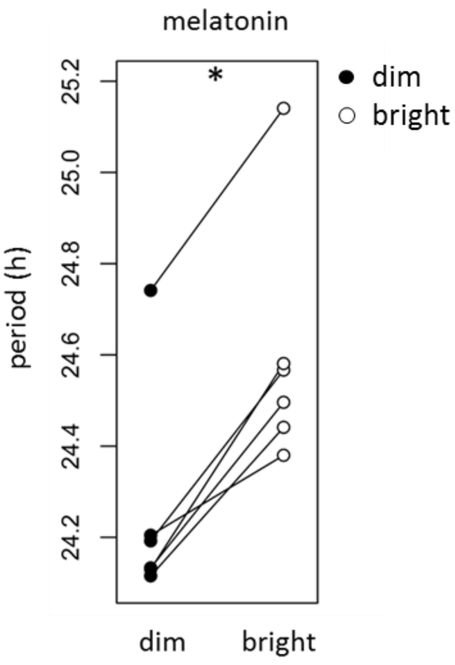


**Figure S4: Internal period ($\tau$) in DL and BL based on melatonin**. Black and white dots represent individual data points collected in DL and BL respectively. Average increase of circadian period under BL versus DL was 21 min.

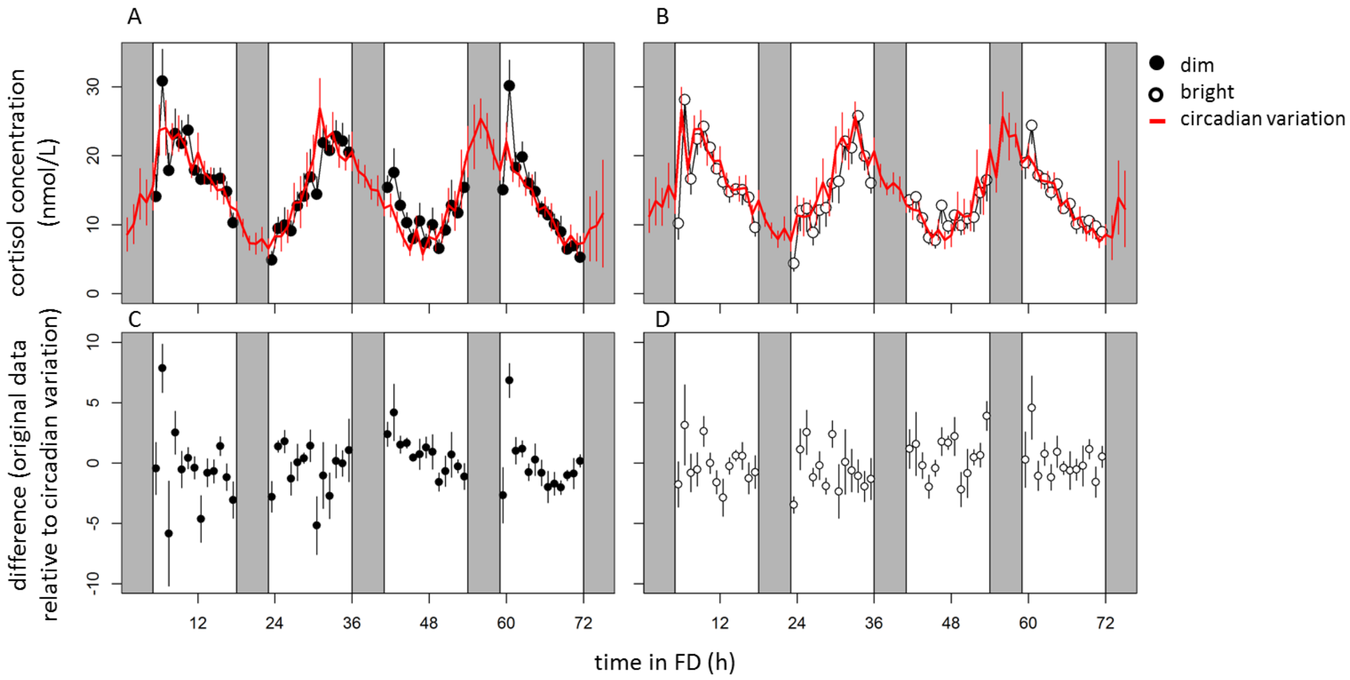


**Figure S5: Stable free run with uniform phase progression in cortisol levels.** Original cortisol concentrations collected during dim (A) and bright light exposure (B) demonstrating their respective circadian variations. After subtracting the calculated circadian variation from the data, the residuals were plotted against time in FD for both DL (C) and BL (D). The residual data (C-D) did not depict a significant circadian not 72-h modulation (p>0.05). The residual variation was 1.61 (DL) and 1.83% (BL) of total variation in the raw data.

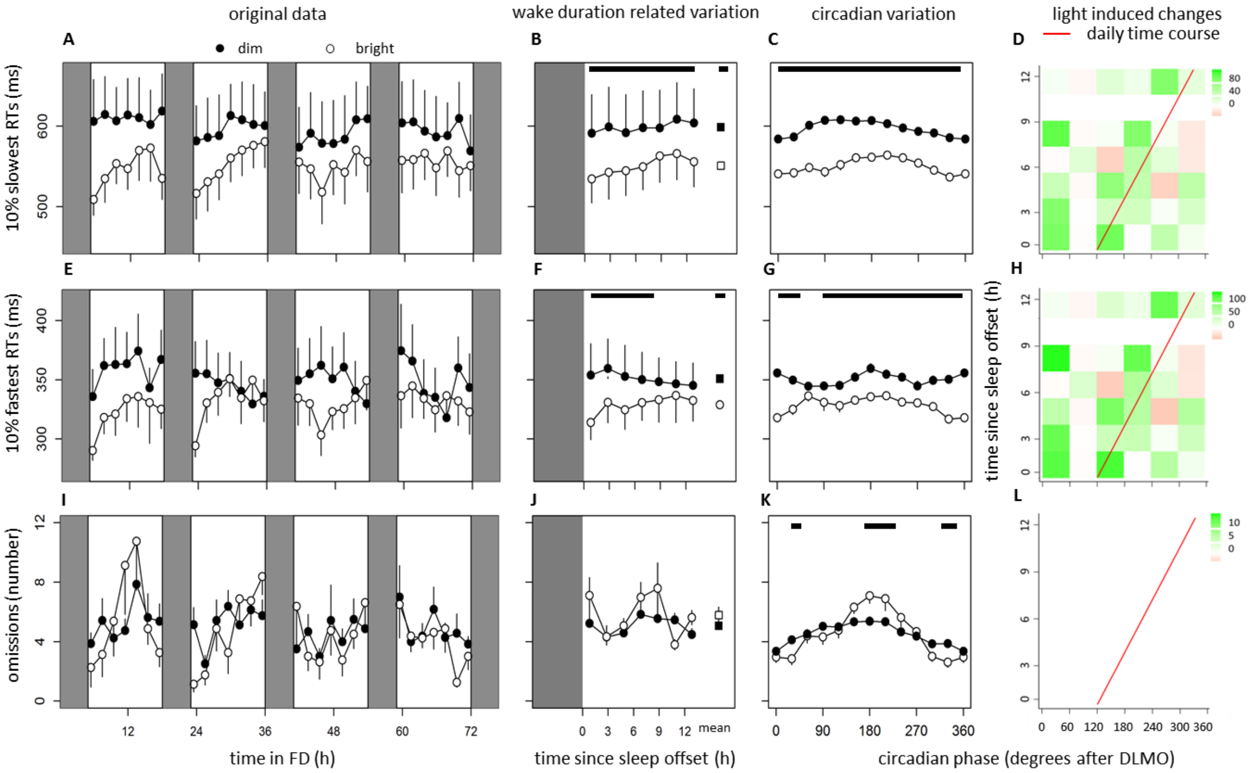


**Figure S6: Data of 10% slowest (top panels), and fastest reaction times (middle panels), and number of omissions (bottom panels).** Time course of 10% slowest (A), and fastest reaction times (E) and number of omissions (I) on the PVT task during the FD protocol. Data replotted as time since sleep offset (B, F and J) and circadian phase in degrees after DLMO (C, G and K), for 10% slowest, and fastest reaction times and number of omission respectively. Contrast analysis describing light induced decrease for all combinations of circadian clock phase and time since sleep offset for 10% slowest (D), and fastest reaction times (H) and number of omissions (L). Data represent mean ± standard error of the mean, with 7 subjects per group. Black dots indicate data collected in dim light, white dots represent data collected in bright light and black and white squares represent averages over all data points under DL and BL respectively. Red line indicates the expected time course over a regular day. Shaded areas represent scheduled sleep (at 0 lux). Significant differences between light conditions (p<0.05) are indicated by horizontal black bars (B,C, F, G, J, K) or colored rectangles (D, H, L).

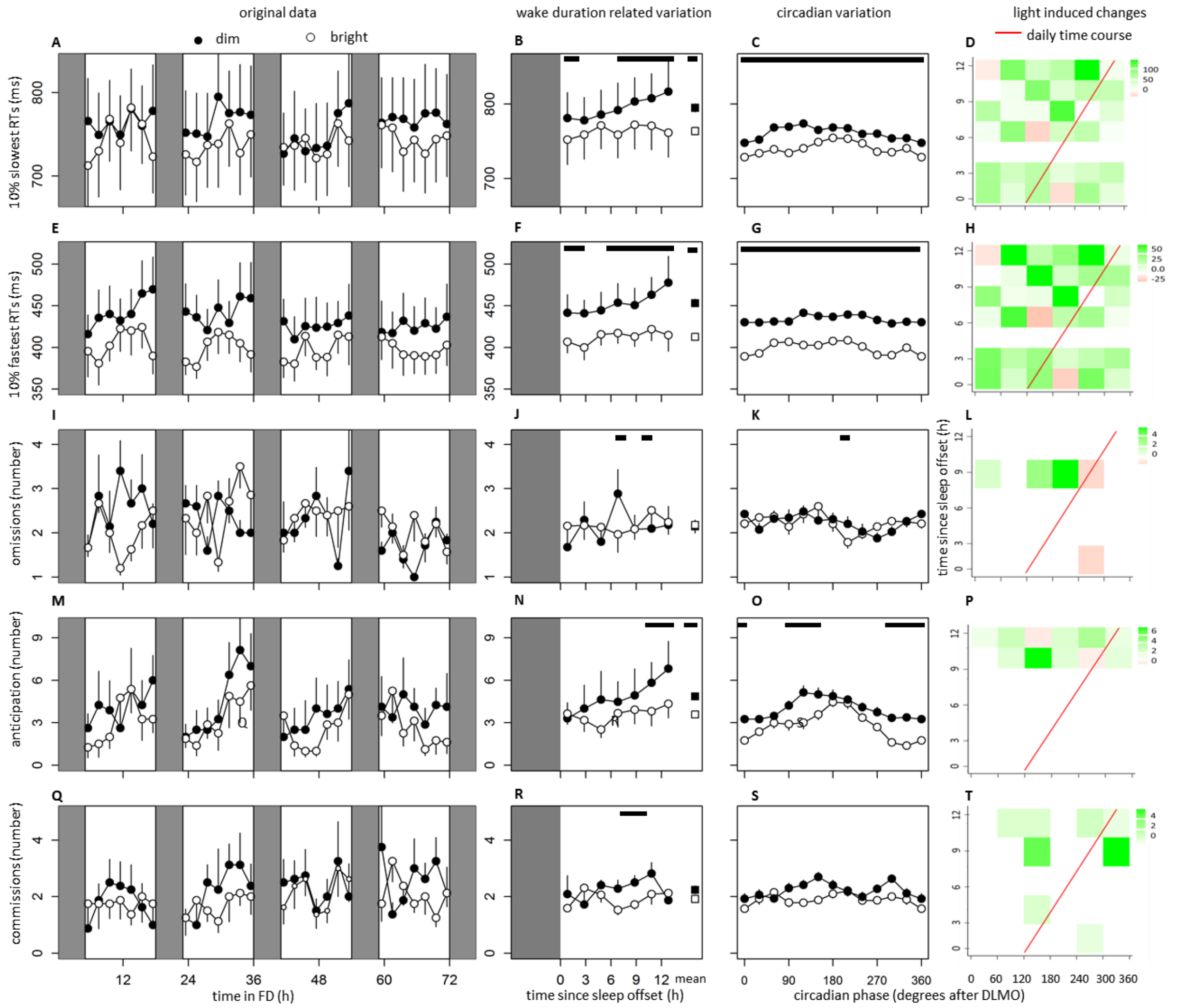


**Figure S7: Data of 10% slowest reaction times (top panels), 10% fastest reaction times, number of omissions and anticipation errors (middle panels) and number of commissions (bottom panels).** Time course of 10%slowest (A) and fastest reaction times (E), as well as errors of omissions (I), anticipation (M) and commission (Q) during the FD protocol. Data replotted as time since sleep offset (B, F, J, N and R) and circadian phase in degrees after DLMO (C, G, K, O and S), for 10%slowest and fastest reaction times, errors of omission, anticipation and commission respectively. Contrast analysis describing light induced decrease for all combinations of circadian clock phase and time since sleep offset for 10% slowest (D) and fastest reaction times (H), as well as errors of omission (L), anticipation (P) and commissions (Q). Data represent mean ± standard error of the mean, with 7 subjects per group. Black dots indicate data collected in dim light, white dots represent data collected in bright light and black and white squares represent averages over all data points under DL and BL respectively. Red line indicates the expected time course over a regular day. Shaded areas represent scheduled sleep (at 0 lux). Significant differences between light conditions ($p<0.05$) are indicated by horizontal black bars (B,C, F, G, J, K, N, O,R,S) or colored rectangles (D, H, L, P, T).

**Table S2: Summary of statistics of sleep-wake related variation (process S), circadian variation (process C), interaction between process S and C, and additive effects of bright light exposure.** Values from linear mixed models on 10% slowest and fastest reaction times, errors of omissions, commissions and anticipation.

| | | **Wake duration related variation (process S)** | | **Circadian variation (process C)** | | **Interaction (process S x C)** | | **Additive effect of bright light** | |
|---|---|---|---|---|---|---|---|---|---|
| **PVT** | **10% slowest** | $F_{(6, 392)}, p$ | 1.43, >0.05 | $F_{(5, 392)}, p$ | 0.00, >0.05 | $F_{(30, 392)}, p$ | 0.00, >0.05 | $F_{(1, 392)}, p$ | **114.32, <0.00001** |
| | **10% fastest** | $F_{(6, 392)}, p$ | 1.63, p>0.05 | $F_{(5, 392)}, p$ | 0.00, >0.05 | $F_{(30, 392)}, p$ | 0.00, >0.05 | $F_{(1, 392)}, p$ | **84.35, <0.00001** |
| | **Omissions** | $F_{(6, 392)}, p$ | **8.47, <0.0001** | $F_{(5, 392)}, p$ | 0.00, >0.05 | $F_{(30, 392)}, p$ | 0.00, >0.05 | $F_{(1, 392)}, p$ | 0.40, >0.05 |
| **GNG** | **10% slowest** | $F_{(6, 392)}, p$ | 1.64, >0.05 | $F_{(5, 392)}, p$ | 0.00, >0.05 | $F_{(30, 392)}, p$ | 0.00, >0.05 | $F_{(1, 392)}, p$ | **63.48, <0.00001** |
| | **10% fastest** | $F_{(6, 392)}, p$ | **2.12, <0.05** | $F_{(5, 392)}, p$ | 0.00, >0.05 | $F_{(30, 392)}, p$ | 0.00, >0.05 | $F_{(1, 392)}, p$ | **38.30, <0.00001** |
| | **Omissions** | $F_{(6, 392)}, p$ | 0.00, >0.05 | $F_{(5, 392)}, p$ | 0.00, >0.05 | $F_{(30, 392)}, p$ | 0.00, >0.05 | $F_{(1, 392)}, p$ | 0.00, >0.05 |
| | **Anticipation** | $F_{(6, 392)}, p$ | **17.62, <0.001** | $F_{(5, 392)}, p$ | 0.00, >.05 | $F_{(30, 392)}, p$ | 0.00, >0.05 | $F_{(1, 392)}, p$ | **21.16, <0.0001** |
| | **Commissions** | $F_{(6, 392)}, p$ | 0.60, >0.05 | $F_{(5, 392)}, p$ | 0.00, >0.05 | $F_{(30, 392)}, p$ | 0.00, >0.05 | $F_{(1, 392)}, p$ | 0.00, >0.05 |

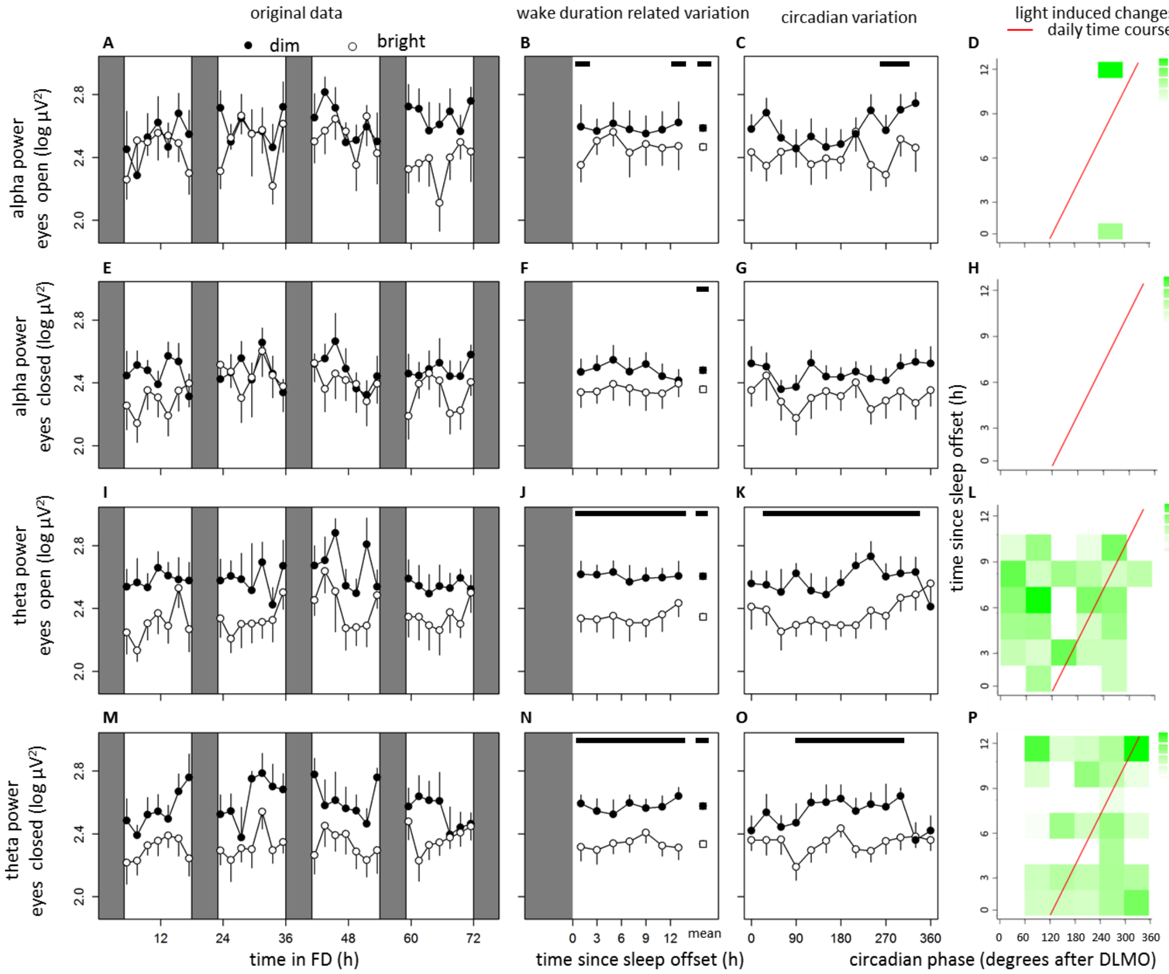


**Figure S8: Data of alpha power with eyes open and closed (top panels) and theta power with eyes open and closed (bottom panels) measured in the frontal electrodes.** Time course of alpha power with eyes open (A), and closed (E), as well as theta power with eyes open (I) and closed (M) during the FD protocol. Data replotted as time since sleep offset (B, F, J, and N) and circadian phase in degrees after DLMO (C, G, K and O), for alpha power with eyes open and closed, as well as theta power with eyes open and closed respectively. Contrast analysis describing light induced decrease for all combinations of circadian clock phase and time since sleep offset for alpha power with eyes open (D), and closed (H) as well as theta power with eyes open (L), and closed (P). Data represent mean ± standard error of the mean, with 7 subjects per group. Black dots indicate data collected in dim light, white dots represent data collected in bright light and black and white squares represent averages over all data points under DL and BL respectively. Red line indicates the expected time course over a regular day. Shaded areas represent scheduled sleep (at 0 lux). Significant differences between light conditions ($p<0.05$) are indicated by horizontal black bars (B,C, F, G, J, K, N, O) or colored rectangles (D, H, L, P).

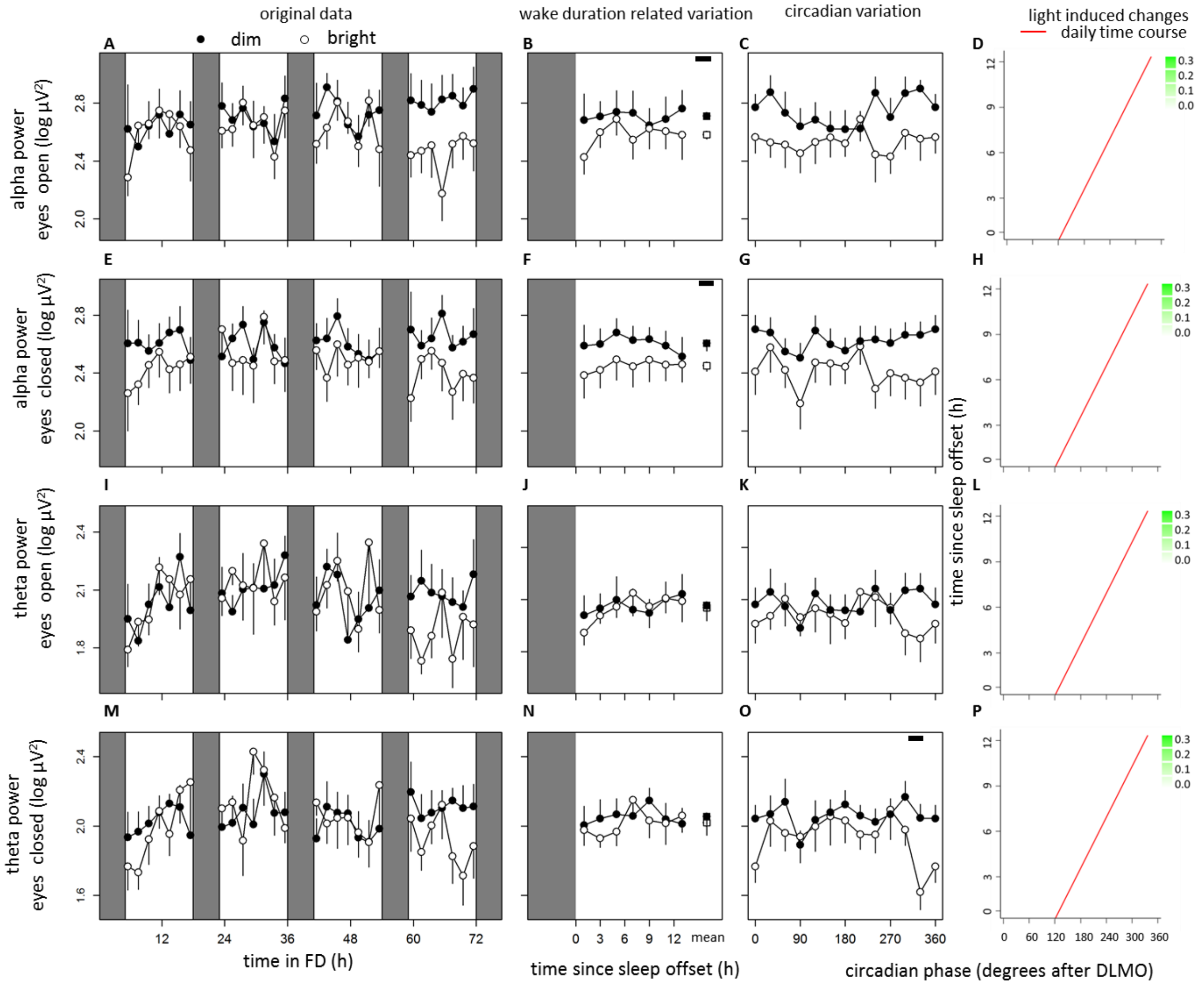


**Figure S9: Data of alpha power with eyes open and closed (top panels) and theta power with eyes open and closed (bottom panels) measured in the occipital electrodes.** Time course of alpha power with eyes open (A), and closed (E), as well as theta power with eyes open (I) and closed (M) during the FD protocol. Data replotted as time since sleep offset (B, F, J, and N) and circadian phase in degrees after DLMO (C, G, K and O), for alpha power with eyes open and closed, as well as theta power with eyes open and closed respectively. Contrast analysis describing light induced decrease for all combinations of circadian clock phase and time since sleep offset for alpha power with eyes open (D), and closed (H) as well as theta power with eyes open (L), and closed (P). Data represent mean ± standard error of the mean, with 7 subjects per group. Black dots indicate data collected in dim light, white dots represent data collected in bright light and black and white squares represent averages over all data points under DL and BL respectively. Red line indicates the expected time course over a regular day. Shaded areas represent scheduled sleep (at 0 lux). Significant differences between light conditions ($p<0.05$) are indicated by horizontal black bars (B,C, F, G, J, K, N, O) or colored rectangles (D, H, L, P).

**Table S3: Summary of statistics of wake duration related variation (process S), circadian variation (process C), interaction between process S and C, and additive effects of bright light exposure.** Values from linear mixed models on EEG based indices of alertness, such as alpha activity with eyes open and closed, and theta activity with eyes open closed in frontal and occipital electrodes.

| | | Wake duration related variation (process S) | | Circadian variation (process C) | | Interaction (process C x S) | | Additive effect of bright light | |
|---|---|---|---|---|---|---|---|---|---|
| **Frontal** | **Alpha eyes open** | $F_{(6, 392)}$, $p$ | 0.53, >0.05 | $F_{(5, 392)}$, $p$ | 1.78, >0.05 | $F_{(30, 392)}$, $p$ | **1.61, <0.05** | $F_{(1, 392)}$, $p$ | **8.25, <0.01** |
| | **Alpha eyes closed** | $F_{(6, 392)}$, $p$ | 0.43, >0.05 | $F_{(5, 392)}$, $p$ | 1.78, >0.05 | $F_{(30, 392)}$, $p$ | 2.75, <0.0001 | $F_{(1, 392)}$, $p$ | **11.18, <0.001** |
| | **Theta eyes open** | $F_{(6, 392)}$, $p$ | 0.53, >0.05 | $F_{(5, 392)}$, $p$ | 1.98, >0.05 | $F_{(30, 392)}$, $p$ | 0.93, >0.05 | $F_{(1, 392)}$, $p$ | **55.47, <0.00001** |
| | **Theta eyes closed** | $F_{(6, 392)}$, $p$ | 0.45, >0.05 | $F_{(5, 392)}$, $p$ | 1.45, >0.05 | $F_{(30, 392)}$, $p$ | 0.80, >0.05 | $F_{(1, 392)}$, $p$ | **56.34, <0.00001** |
| **Occipital** | **Alpha eyes open** | $F_{(6, 392)}$, $p$ | 0.56, >0.05 | $F_{(5, 392)}$, $p$ | 1.37, >0.05 | $F_{(30, 392)}$, $p$ | 0.97, >0.05 | $F_{(1, 392)}$, $p$ | **7.42, <0.01** |
| | **Alpha eyes closed** | $F_{(6, 392)}$, $p$ | 0.56, >0.05 | $F_{(5, 392)}$, $p$ | 0.70, >0.05 | $F_{(30, 392)}$, $p$ | 0.99, >0.05 | $F_{(1, 392)}$, $p$ | **11.29, <0.001** |
| | **Theta eyes open** | $F_{(6, 392)}$, $p$ | 0.38, >0.05 | $F_{(5, 392)}$, $p$ | 0.24, >0.05 | $F_{(30, 392)}$, $p$ | 1.05, >0.05 | $F_{(1, 392)}$, $p$ | 0.24, >0.05 |
| | **Theta eyes closed** | $F_{(6, 392)}$, $p$ | 0.91, >0.05 | $F_{(5, 392)}$, $p$ | 1.14, >0.05 | $F_{(30, 392)}$, $p$ | 1.22, >0.05 | $F_{(1, 392)}$, $p$ | 0.70, >0.05 |

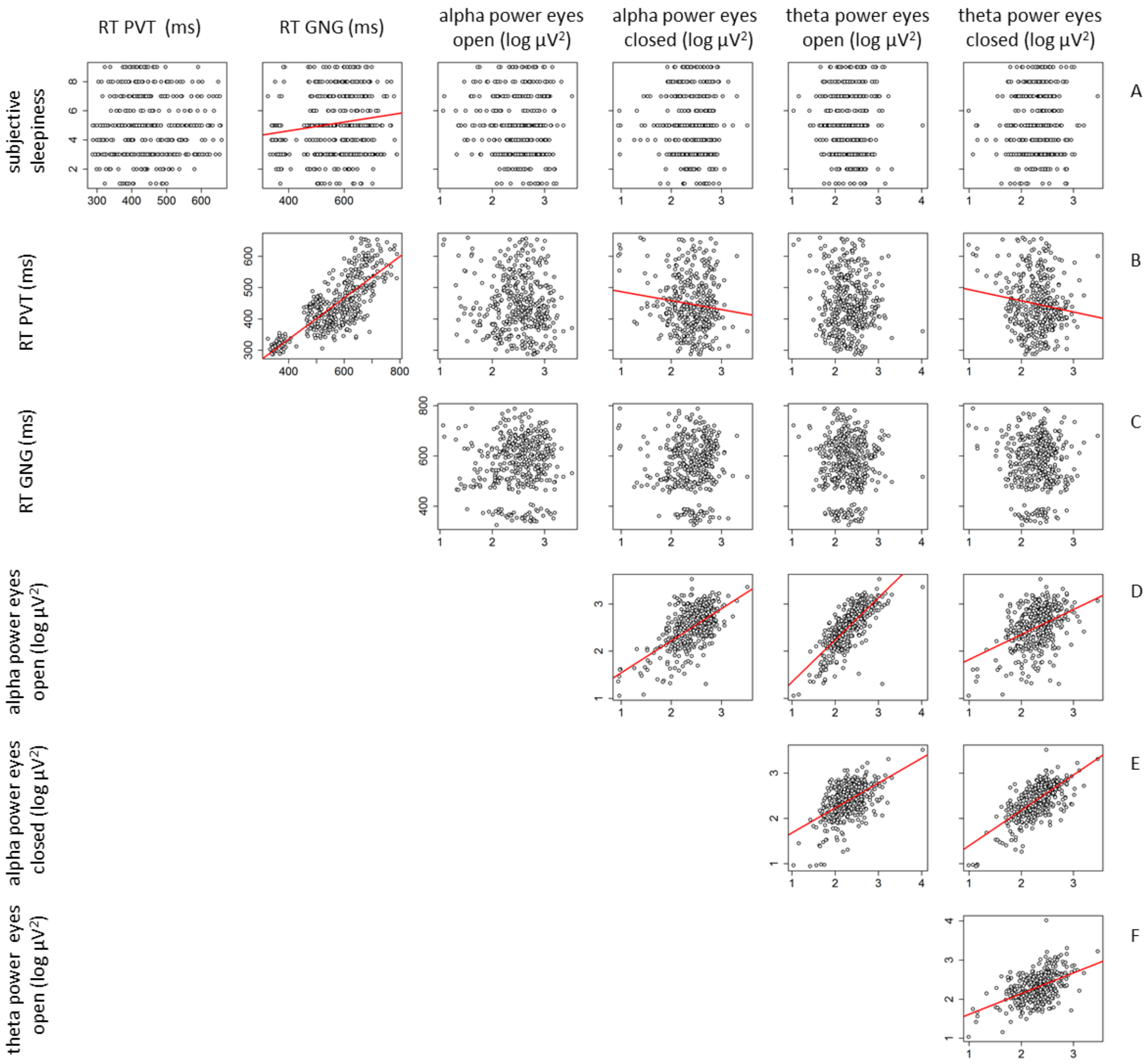


**Figure S10: Correlation matrix of parameters of alertness.** Every dot represents one data point. Depicted are the correlation between subjective alertness and reaction time on the PVT, Go-NoGo and parameters of EEG (A), reaction time on the PVT, GNG, and EEG parameters (panel B), GNG reaction time and EEG parameters (C), and of EEG measures (D-F). Significant correlations ($p<0.05$) are indicated by a red line.
